# Profiling and Quantification of Anthocyanins in Novel Purple-Pericarp Sweetcorn and Purple-Pericarp Maize

**DOI:** 10.1101/2022.07.25.501484

**Authors:** Apurba Anirban, Hung T. Hong, Tim J O’Hare

**Affiliations:** Centre for Nutrition and Food Sciences, Queensland Alliance for Agriculture & Food Innovation (QAAFI), The University of Queensland, Brisbane, Australia

**Keywords:** anthocyanin, cyanidin, starchy maize, purple sweetcorn, white sweetcorn

## Abstract

Anthocyanins, secondary metabolites of pigmented corns consisting of cyanidin-, pelargonidin- and peonidin-based glucoside. While cyanidin-peonidin types are responsible for purple-pigmentation, pelargonidin type is responsible for red-pigmentation. This study examined anthocyanin concentrations in a novel purple-pericarp *shrunken2* sweetcorn ‘Tim’ lines in comparison to the parental purple-pericarp ‘Costa Rica’ maize’ and white sweetcorn ‘Tims-white’ lines. The study found similar concentrations of anthocyanin in both ‘Tim1’ and the ‘Costa Rica’, at sweetcorn eating stage, whereas ‘Tims-white’ has no detectable anthocyanin. Total anthocyanins found in the ‘Costa Rica’ and ‘Tim1’, ‘Tim2’, ‘Tim4’ and ‘Tim5’ lines were 255.79, 253.03, 238.10, 198.66, and 221.36 mg/100 g FW (fresh weight), respectively. In all the developed purple-sweetcorn ‘Tim’ lines along with the purple maize parent, the cyanidin-peonidin (purple) proportion of total anthocyanin was 78-93%, as the predominant anthocyanin pigment. The anthocyanin concentration in ‘Tim1’ at eating stage was significantly much higher than currently exists with other coloured fruits.

## 1. Introduction

Purple *shrunken2* (*sh2*) sweetcorn is uncommon due to a tight genetic linkage between *sh2* and *a1* (*anthocyaninless1*) genes (Yao et al., 2002). In purple maize (*A1Sh2.A1Sh2*), the anthocyanin biosynthesis dominant structural gene *anthocyanin1* (*A1*) is closely linked to the starch biosynthesis dominant structural gene, *Sh2* (Kramer et al., 2015). As a result, the kernels are purple and round, due to the presence of starch, rather than sugar, when fully mature. In contrast, in yellow/white sweetcorn, the recessive *a1* gene is closely linked to the recessive *sh2* gene (Civardi et al., 1994). Therefore, the kernels are non-purple and shrunken, due to the absence of starch, when fully mature. Recently, a novel purple *sh2* sweetcorn was developed from Peruvian purple maize (‘Costa Rica’) and white *sh2* sweetcorn (‘Tims-white’) by breaking the extremely tight genetic link between the non-purple *a1* and supersweet *sh2* genes (Anirban and O’Hare, 2020, Anirban et al., 2022).

The appearance of the purple colour is due to the synthesis of anthocyanins in the kernel pericarp tissue or/and in the kernel aleurone. (Emerson, 1921), while sweetcorn lacks any anthocyanin pigmentation. In pericarp-coloured maize the anthocyanin concentration is eight-fold higher than that found in aleurone-coloured maize (Paulsmeyer et al., 2017). Anthocyanins previously identified in purple non-sweet maize and purple *brittle1* sweetcorn are derivatives of anthocyanin, *e.g.*, cyanidin, peonidin and pelargonidin (Styles and Ceska, 1972), and their malonylated (Pascual-Teresa et al., 2002) as well as dimalonylated (Hong et al., 2020) derivatives.

Research by Wu et al., (2017) conducted on obese mice revealed that anthocyanin lowered the body weight as well as risk of colon cancer being reduced. Study by Warner et al., (2014) found freeze-dried black raspberries prevented oral cancer development in humans. Another study by Miranda-Rottmann et al., (2002) found anthocyanins in berry juice with higher antioxidant capacity inhibited LDL (low-density lipoprotein) oxidation. Anthocyanin-rich fresh juices from strawberry fruit significantly inhibited mutagenesis caused by methyl methane sulfonate (MMS) responsible for producing cancer in humans (Hope Smith et al., 2004). Review by Lao et al., (2017) indicated that Japanese-, Chinese- and Peruvian-purple maize anthocyanins act against colon-, prostrate-, mammary-, liver- and kidney-cancer of different rats and mice as well as prevented hypertension in humans. In male rats, purple maize natural anthocyanin inhibited colorectal carcinogenesis (Hagiwara et al., 2001). Therefore, anthocyanin found in the developed purple *shrunken2* sweetcorn could reduce different diseases. Sweetcorn is harvested and eaten when the kernel is still physiologically immature (Khanduri et al., 2011), in contrast to purple maize that is used for flour or food colouring (Lago et al., 2014). Therefore, the differences in anthocyanin concentrations and profiles of them need to investigate.

Anthocyanin levels in the purple sweetcorn kernel increase with increasing kernel maturity (Hong et al., 2020). Anthocyanin development begins at the point of silk attachment, gradually covering the surface of the kernel as the kernel matures (Hong et al., 2021). However, the anthocyanin profile of novel purple-pericarp *sh2* sweetcorn has not previously been analysed, either at sweetcorn eating stage or at its full-kernel maturity. It is expected that anthocyanin profile of the progenies should have similarity with its ‘Costa Rica’ purple maize parent. As such, the primary aim of current study was to document anthocyanin concentrations of these different lines, and how these anthocyanin profiles compare to the parent lines, at both sweetcorn eating stage and at full kernel maturity.

## 2. Materials and methods

### 2.1 Plant materials

Cobs from the purple-pericarp Peruvian maize parental line, ‘Costa Rica’, white sweetcorn parental lines, ‘Tims-white’, and their F3 and F6 progenies (Fig 1) were investigated in this current study. ‘Costa Rica’ is a purple-pericarp starchy maize (*A1Sh2.A1Sh2*), ‘Tims-white’ is a *shrunken2* (*sh2*) white sweetcorn (*a1sh2.a1sh2*), F3 is the developed heterozygous purple-pericarp *sh2* sweetcorn line (*A1a1.sh2sh2*), and F6 line ‘Tim1’ is the homozygous purple-pericarp *sh2* sweetcorn line (*A1A1.sh2sh2*), while the remainder of the F6 lines are still heterozygous (*A1a1.sh2sh2*), *i.e.*, segregating for the *A1* gene.

**Fig 1.**
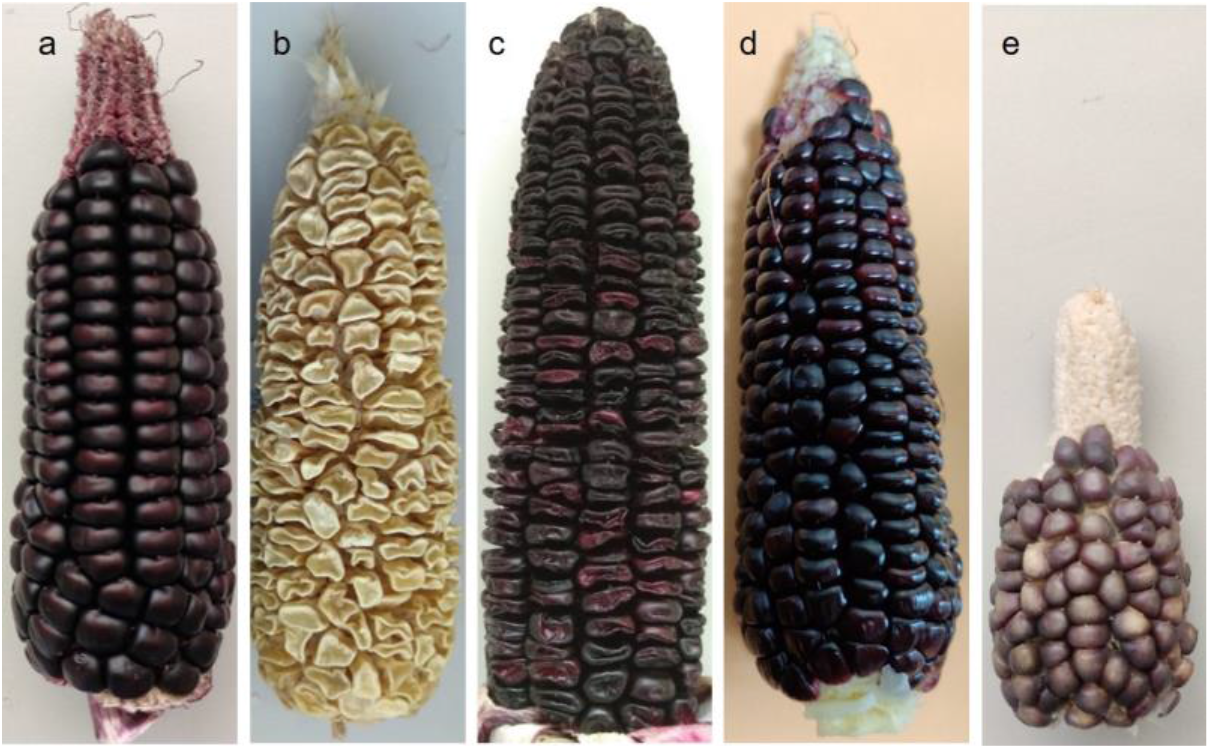
Cobs of parental lines, F3 and F6 progenies, as well as aleurone-pigmented corn: **a)** purple-pericarp maize, ‘Costa Rica’ (*A1Sh2.A1Sh2*), **b)** white sweetcorn, ‘Tims-white’ (*a1sh2.a1sh2*), **c)** F3 heterozygous purple-pericarp sweetcorn (*A1a1.sh2sh2*), **d)** F6 homozygous purple-pericarp sweetcorn (*A1A1.sh2sh2*), and **e)** aleurone-pigmented (non-pericarp pigmented) corn (‘Tims-aleurone’).

Initially, three heterozygous F3 purple-pericarp sweetcorn lines (‘Tim1’, ‘Tim2’, and ‘Tim3’) were obtained, and then two other lines were detected (‘Tim4’ and ‘Tim5’). Therefore, there were five segregants displaying purple-pericarp *sh2* traits at the F3 stage (‘Tim1’ to ‘Tim5’), however one of them (‘Tim3’) was discarded due to *Fusarium* infection. Purple aleurone kernels (‘Tims-aleurone’), not related to the purple-pericarp parent or progeny, were also used for comparison of aleurone anthocyanin profile with pericarp anthocyanin profile. Mature seeds of 60 DAP (days after pollination) and sweetcorn eating stage kernels of 25 DAP were used for the study. Kernels of parental lines (‘Costa Rica’ and ‘Tims-white’), aleurone-pigmented kernels (‘Tims-aleurone’) and F3 progenies at the mature stage were assessed for anthocyanin profile and concentration, whereas parental lines and the F6 purple-pericarp sweetcorn lines at the sweetcorn eating stage (25 DAP) were compared.

### 2.2 Anthocyanin extraction, identification and quantification

#### 2.2.1 Chemicals

Standards for Cy3G (Cyanidin-3-glucoside), Pg3G (pelargonidin-3-glucoside) and Pn3G (peonidin-3-glucoside) were received from Extrasynthese (Genay, France). Solvents and other chemicals (HPLC or analytical grade) were procured from Sigma-Aldrich (Sydney, NSW, Australia). Deionized pure water (Millipore Australia Pty Ltd, Kilsyth, VIC, Australia) was utilized in this research.

#### 2.2.2 Anthocyanin extraction

Approximately 100 kernels were randomly separated from different locations on the cob and immediately snap frozen using liquid nitrogen. Subsequently, the samples were stored at -20 °C approximately up to three months prior to analysis. Approximately 15 kernels were put into milling vessels and placed in liquid nitrogen for 3 min. Then the vessels were transferred into a MM400 Retsch Mixer Mill (Haan, Germany) which was operated at 30 Hz for 60 sec. Frozen powdered kernel (subsample, 0.5 g) was accurately weighed and used for the estimation of individual anthocyanin concentration. Anthocyanins extraction from the frozen powder sample was performed according to a previous study by Hong et al. (2020). Briefly, frozen powdered sample (about 0.5 g) was placed into a falcon tube (50 mL) and added extraction solution (aqueous 80% methanol, 0.1 M HCl) of 3 mL (cold) at 4 °C. reciprocating shaker (horizontal) RP 1812 (Victor Harbor, SA, Australia) was used to shake the mixer at 250 rpm for 10 min under 4 °C cool temperature and dim light and sonicated at 4 °C for 10 min. Centrifugation was performed at 4 °C, at 4000 rpm for 10 min (Eppendorf Centrifuge 5804, Germany). The supernatant of the samples was collected, and the pellet reextracted twice following the similar process. Combined supernatants were filtered through a 0.20 μm hydrophilic PTFE syringe filter into HPLC vials to perform chemical analysis. The extraction was performed in triplicate.

#### 2.2.3 Anthocyanin identification

Anthocyanin profiles were obtained using UHPLC-DAD-MS (ultra-high performance liquid chromatography–diode array detection-mass spectrometry) (Shimadzu, Kyoto, Japan) as described previously by Hong et al., (2021). For operation of the instrument and data-processing LCMS software (Lab solutions) (Ver.5.85; Shimadzu) was used. Chromatographic separation was performed on a reverse phase Acquity UPLC BEH C18 column (100 × 2.1 mm i.d., 1.7 particle size; Waters, Dublin, Ireland). Temperature of column was maintained at 50 °C and scanning of DAD spectrum was performed from 200 to 600 nm. The elution was performed with 92% water, 7% acetonitrile and 1% formic acid (100% of mobile phase A) as an initial isocratic hold for 1 min, followed by a linear gradient from 100% to 85% of mobile phase A for 30 min, purging for 3 min at 100% of mobile phase B (acetonitrile, 1% formic acid), conditioning for 1 min, and re-equilibration for 5 min.

#### 2.2.4 Anthocyanin quantification

Anthocyanins were quantified using a UHPLC–DAD Agilent 1290 Infinity system (Agilent Technologies, USA; System 2). The UHPLC consisted of a diode-array (DAD) detector (1290 Infinity), pump (1290 Infinity), column oven (1290 Infinity), auto-sampler (1290 Infinity) and a system controller (1290 Infinity). Agilent HassHunter workstation Data Acquisition version B.07.00/Build 7.0.7022.0 was controlled by the UHPLC-DAD system. Agilent MassHunter Quantitative Analysis Version B.07.00/Build 7.0.457.0 was employed for data analysis. Chromatographic separation was achieved on a reversed-phase Acquity UPLC BEH C18 column (150 × 2.1 mm i.d., 1.7 μm particle size; Waters, Dublin, Ireland) at 50 °C with a flow rate of 0.25 mL/min. The DAD spectra were scanned from 200 to 800 nm and anthocyanins were detected at 520 nm. The concentration of individual anthocyanins was determined by calibration curves (external) of Cy3G, Pn3G and Pg3G, and their respective malonated counterparts. To define purple colour, cyanidin and peonidin were pooled together, as peonidin is a methylated form of cyanidin (Petrussa et al., 2013).

### 2.3 Objective colour measurement

Mature seeds from the round (non-shrunken) ‘Costa Rica’ and F3 purple shrunken progenies (‘Tim1’, ‘Tim2’, ‘Tim4’ and ‘Tim5’) were used for the study to identify the impact of total anthocyanin on kernel pericarp colour. Colour was measured objectively (McGuire, 1992) for C* (chroma) and H* (hue angle) with a Chroma meter, Minolta CR-400 (Konica Minolta, Osaka, Japan). Ten kernels were removed from the cob and positioned upright on a black plasticine base to replicate their original placement on the cob.

### 2.4 Statistical analysis

A statistical software known as Prism (GraphPad Prism 9.3.1) was used for one-way factorial analysis of variance (ANOVA) to assess variances and pairwise multiple comparisons. To compare differences between means, Fisher’s Least significant difference (LSD, p < 0.005) was used. Three replicates (15 kernels for each sample) were used from a cob for statistical analysis.

### 2.5 Kernel anatomy

To observe whether anthocyanin is developed in the pericarp (outer layers), aleurone layer (inner single layer, from pericarp to endosperm), and endosperm, a longitudinal section (LS) of mature kernels (60 DAP) of the parents and progenies were assessed.

### 2.6 Kernel Physiology

To understand the stage of kernel development on anthocyanin development, developing kernels were visually inspected from seven to 28 DAP in the field. Photographs were taken to visualize anthocyanin development across the kernel.

## 3. Results

### 3.1 Anthocyanin quantification of mature round (starchy) kernels

Cyanidin, peonidin and pelargonidin based anthocyanin compounds at 60 DAP (day after pollination) were identified by UHPLC-DAD-MS. The total anthocyanin concentration of mature kernels of ‘Costa Rica’ was significantly much higher (194.47 mg/100g FW) than the ‘Tims-aleurone’ maize genotype (27.65 mg/100g FW) (Fig 2a). By contrast, there was no anthocyanin detected in the white shrunken ‘Tims-white’ parental line. No significant difference (P<0.005) existed between ‘Costa Rica’ and ‘Tims-aleurone’ regarding pelargonidin concentration, however they were significantly different regarding cyanidin, peonidin and cyanidin with peonidin concentrations (Fig 2a). The anthocyanin profiles of ‘Costa Rica’ consisted of 97% cyanidin (with peonidin) and 3% pelargonidin. By contrast, ‘Tims-aleurone’ genotypes had 90% cyanidin (with peonidin) and 10% pelargonidin.

**Fig 2.**
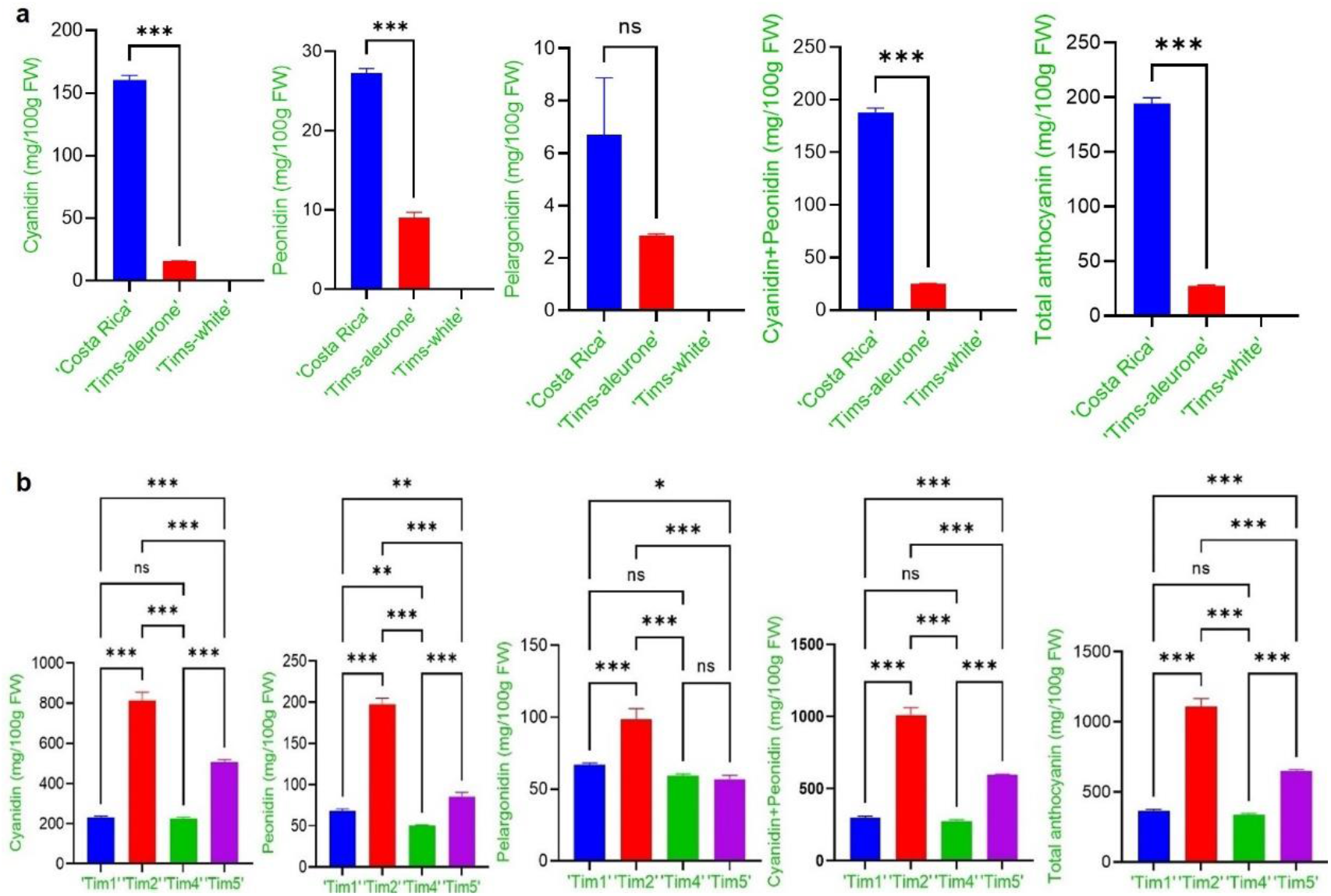
Multiple comparison showing ns=non-significant, *=significant at P < 0.005, **= significant at P <0.002, and ***= significant at P < 0.001 regarding profiles of anthocyanin of mature kernels: **a)** profiles of anthocyanin of mature round (starchy) parental line (‘Costa Rica’) and aleurone pigmented (‘Tims-aleurone’) round kernels at 60 DAP. **b)** profiles of anthocyanin of developed F3 purple-shrunken kernels at 60 DAP.

### 3.2 Quantification of anthocyanin of mature shrunken (non-starchy) kernels

The anthocyanin concentrations of the F3 shrunken segregants (‘Tim1, ‘Tim2’, ‘Tim4’ and ‘Tim5’) at 60 DAP were estimated. The highest total anthocyanin concentration (1109.30 mg/100g FW) was observed in the ‘Tim2’ purple shrunken genotype, which is significantly much higher (P<0.005) than other genotypes (Fig 2b). In comparison, the ‘Tim1’, ‘Tim4’ and ‘Tim5’ genotypes contained 365.35, 335.28 and 649.32 mg/100g FW total anthocyanin, respectively, which were significantly different (P<0.005), except ‘Tim1’ and ‘Tim4’ lines. There were significant differences (P<0.005) regarding cyanidin, peonidin, cyanidin with peonidin and pelargonidin concentrations between the individual ‘Tim’ lines, however there was no significant difference between ‘Tim1’ and ‘Tim4’, except for peonidin (Fig 2b). In addition, the ‘Tim2’ and ‘Tim5’ anthocyanin profiles consisted of 91% cyanidin (with peonidin) and 9% pelargonidin, respectively, whereas ‘Tim1’ and ‘Tim4’ genotypes had 82% and 18%, respectively.

### 3.3 Quantification of anthocyanin derivatives of eating-stage (25 DAP) kernels of parental and F6 ‘Tim’ lines

Analysis of anthocyanin derivatives at the sweetcorn eating stage (25 DAP) identified nine cyanidin-, peonidin- and pelargonidin-based (Fig 3a) anthocyanin derivatives in the purple-pericarp sweetcorn accessions (Fig 3b). These included cyanidin-3-glucoside (Cy3G), cyanidin-3-malonyl glucoside (Cy3MG), cyanidin-3-dimalonyl glucoside (Cy3DMG); peonidin-3-glucoside (Pn3G), peonidin-3-malonyl glucoside (Pn3MG), peonidin-3-dimalonyl glucoside (Pn3DMG), pelargonidin-3-glucoside (Pg3G), pelargonidin-3-malonyl glucoside (Pg3MG), and pelargonidin-3-dimalonyl glucoside (Pg3DMG).

**Fig 3.**
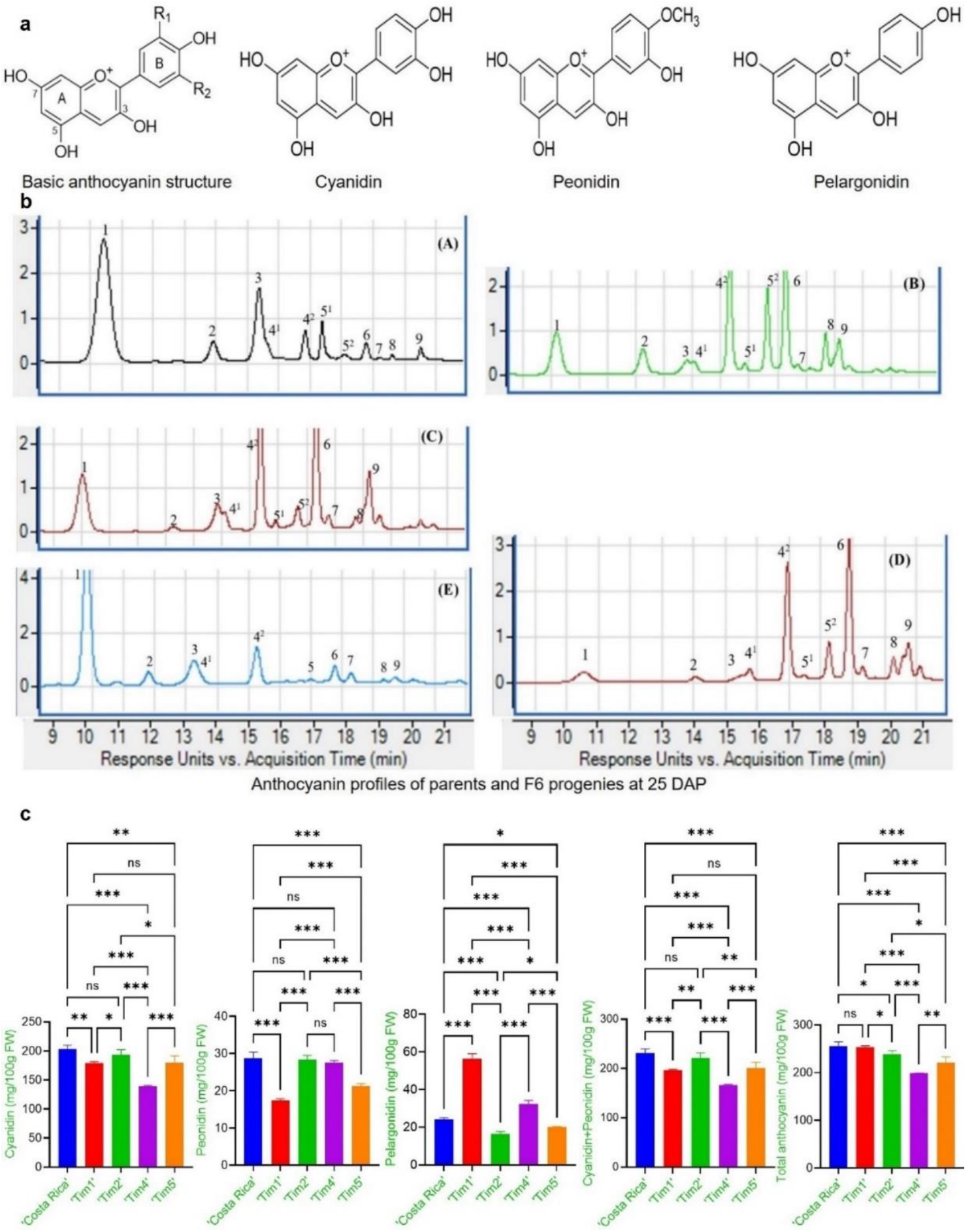
Structure and profiles of anthocyanin of parents and developed F6 lines at 25 days after pollination: **a)** basic anthocyanin structure, as well as structure of cyanidin, peonidin and pelargonidin; **b)** profiles of anthocyanin of ‘Costa Rica’ and different F6 progenies at 25 DAP: (A) ‘Costa Rica’, (B) ‘Tim1’, (C) ‘Tim2’, (D) ‘Tim4’, and (E) ‘Tim5’ obtained from UHPLC-DAD-MS at 520nm, (1): cyanidin-3-glucoside, (2): pelargonidin-3-glucoside, (3): peonidin-3-glucoside, (4^1^): cyanidin-3-malonyl glucoside, (4^2^): cyanidin-3-malonyl glucoside isomer, (5^1^): pelargonidin-3-malonyl glucoside, (5^2^): pelargonidin-3-malonyl glucoside isomer, (6): cyanidin-3-dimalonyl glucoside, (7): peonidin-3-malonyl glucoside, (8): pelargonidin-3-dimalonyl glucoside, (9): peonidin-3-dimalonyl glucoside; **c)** multiple comparison showing ns=non-significant, *=significant at P < 0.005, **= significant at P <0.002, and ***= significant at P < 0.001) regarding profiles of anthocyanin of eating stage kernels.

It was found that the sweet purple-pericarp developed line ‘Tim1’ (F6) produced 253.03 mg/100g FW total anthocyanin in comparison to its purple maize parental line ‘Costa Rica’ (255.79 mg/100g FW), which was not significantly different (P<0.005) (Fig 3c). Again, there was no anthocyanin detected in the white sweetcorn parent, ‘Tims-white’, at 25 DAP. Total anthocyanins found in ‘Tim2’, ‘Tim4’ and ‘Tim5’ were 238.10 mg/100g FW, 198.66 mg/100g FW, and 221.36 mg/100g FW, respectively, which were significantly different (P<0.005) from each other (Fig 3c).

The main anthocyanin component found in all lines was cyanidin. Although there was no significant difference between ‘Tim1’ and ‘Costa Rica’ regarding total anthocyanin concentration at 25 DAP, however, significant differences in cyanidin, peonidin, cyanidin with peonidin, and pelargonidin concentrations existed (Fig 3c). There was no significant difference (P<0.005) between ‘Costa Rica’ and ‘Tim2’, or between ‘Tim1’ and ‘Tim5’ regarding cyanidin with peonidin concentration, whereas all lines were significantly different regarding pelargonidin concentration (Fig 3c).

### 3.4 Objective colour measurement of the mature round (starchy) and shrunken (non-starchy) kernels

The chroma (C*) values of mature round kernels of ‘Costa Rica’, as well as F3 mature shrunken ‘Tim1’, ‘Tim2’, ‘Tim4’ and ‘Tim5’ kernels (Fig 4a) were 5.5, 7.7, 8.1, 6.6 and 8.1, respectively. The C* value of mature round kernels of ‘Costa Rica’ was significantly different (P<0.005) than all F3 mature shrunken kernels, except ‘Tim4’ (Fig 4b). The C* value of ‘Costa Rica’ was also lower than all other shrunken lines, indicating its less-intense colour. No significant (P<0.005) difference was found between other F3 mature shrunken lines regarding C* value, except ‘Tim2’ and ‘Tim4’.

**Fig 4.**
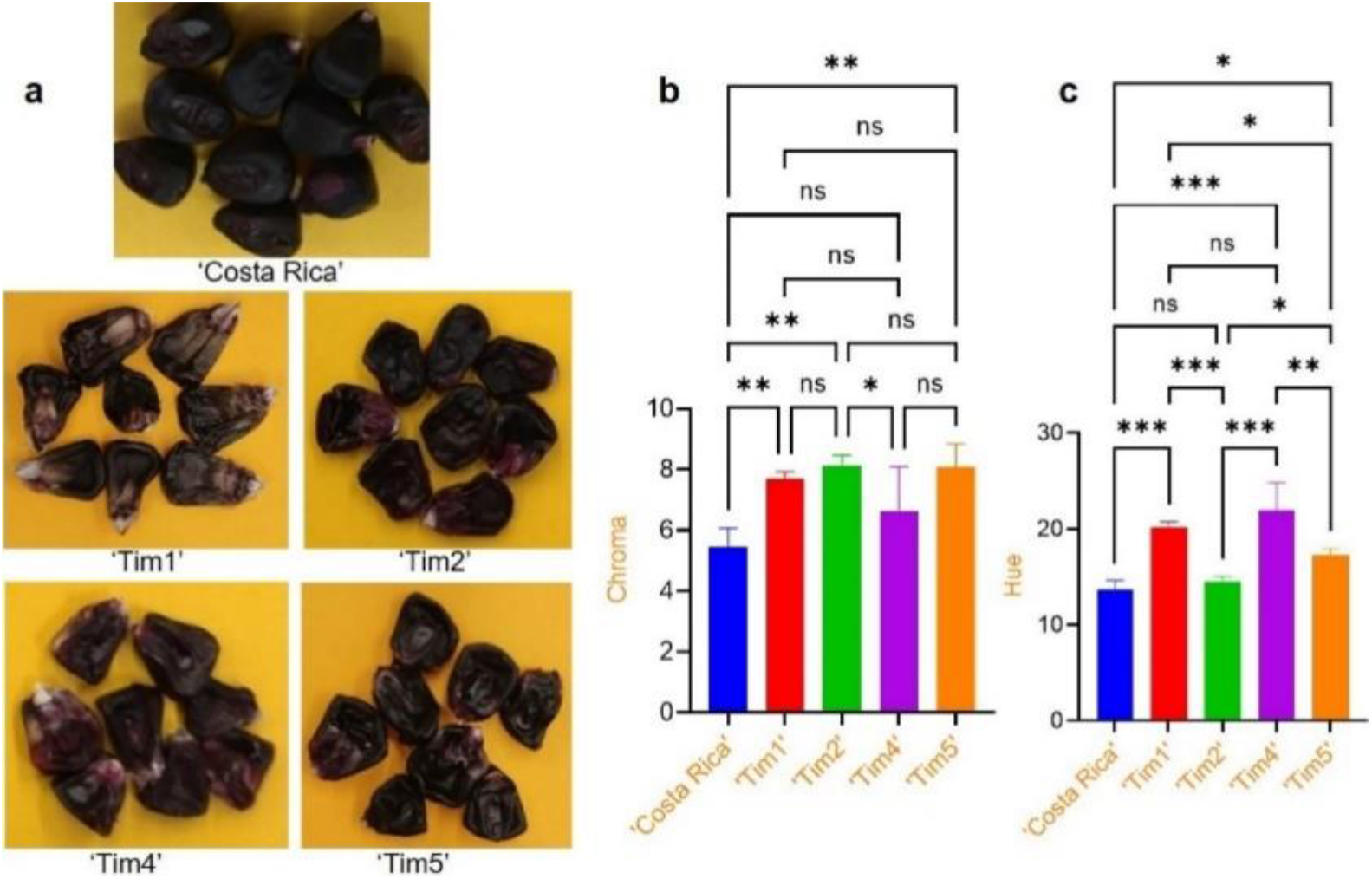
Objective colour measurement of mature kernels: **a)** mature round kernels of ‘Costa Rica’ and F3 purple shrunken kernels (‘Tim1’, ‘Tim2’, ‘Tim4’ and ‘Tim5’); **b)** comparative analysis of chroma values of mature kernels of ‘Costa Rica’ and mature F3 shrunken kernels; **c)** comparative analysis of hue values of mature kernels of ‘Costa Rica’ and mature F3 ‘Tim1’, ‘Tim2’, ‘Tim4’ and ‘Tim5’ shrunken kernels. Multiple comparison indicating ns=non-significant, *= significant at P <0.005, **=significant at P < 0.033, and ***= significant at P < 0.001.

The hue (H*) values of mature round kernels of ‘Costa Rica’ and F3 mature shrunken ‘Tim1’, ‘Tim2’, ‘Tim4’ and ‘Tim5’ kernels were 13.9, 20.22, 14.43, 21.89 and 17.25, respectively. The H* value of ‘Costa Rica’ was significantly different (P<0.005) than all F3 mature shrunken kernels, except ‘Tim2’ (Fig 4b). The H* value of the ‘Tim2’ purple shrunken genotype was lower than all other genotypes, indicating more purple colour in comparison to the other purple shrunken genotypes. Except ‘Tim1’ and ‘Tim4’, significant (P<0.005) differences were present among the lines regarding H* value, which indicate that colour variation also exists among them.

### 3.5 Kernel anatomy regarding anthocyanin development

Anthocyanin development in different tissues of different lines is shown in Fig 5a. The mature ‘Costa Rica’ parental line had signs of anthocyanin pigmentation (purple colour) in the pericarp, aleurone and vitreous endosperm, but not in the starchy endosperm. In contrast, the mature ‘Tims-aleurone’ kernel had anthocyanin only in the aleurone layer and vitreous endosperm (outside pericarp was transparent, which was confirmed by visual inspection), and the ‘Tims-white’ parent had no anthocyanin in any layer.

**Fig 5.**
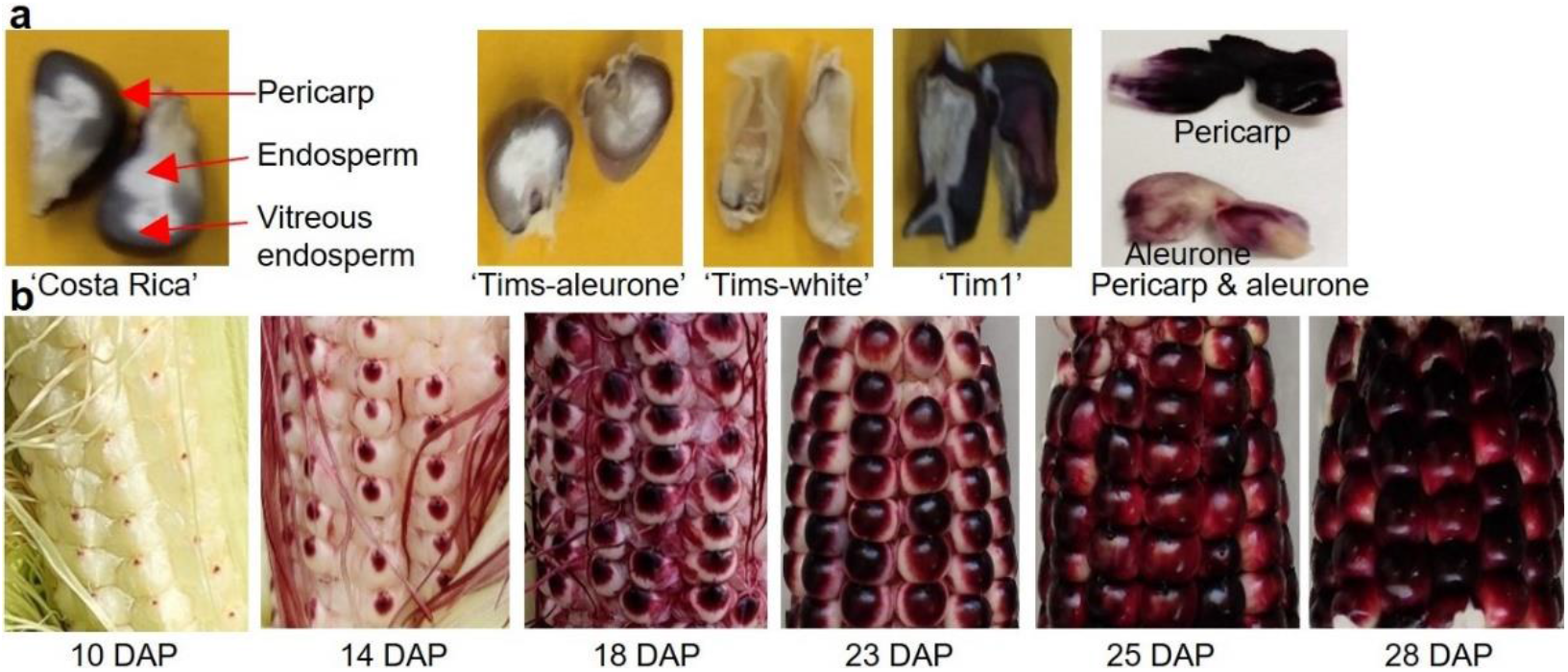
Anatomy of kernels and physiology of anthocyanin development. **a)** LS of different kernels: ‘Costa Rica’, ‘Tims-aleurone’, ‘Tims-white’, F3 line ‘Tim1’; and separated pericarp and aleurone tissues of F3 line ‘Tim1’; **b)** physiology of anthocyanin development at different stages of development (10-28 DAP).

The F3 mature purple shrunken kernels had shrivelled kernels and anthocyanin was confined to the pericarp, aleurone layer, and vitreous endosperm (image of only one shrunken kernel of the F3 line, ‘Tim1’ is shown in Fig 5a). The sections also showed that there was no visual evidence of anthocyanin development in the embryo and starchy endosperm of any of the genotypes assessed. These all mature kernels were used for the quantification of anthocyanin at 60 DAP.

### 3.6 Kernel maturity and anthocyanin development

Visual inspection was performed to observe the stages of anthocyanin development in kernels at different stages of development (Fig 5b). Anthocyanin accumulation was observed to start at the kernel tip (point of attachment of the silk) at 10 days after pollination. Anthocyanin accumulation gradually increased from 10 to 18 DAP and started to spread across the surface of the kernel. At 23 DAP, more than three-quarters of the kernel was purple. By 25 DAP, the kernels at the lower part of the cob were fully purple, and some kernels of the upper cob were still turning purple. Finally, at 28 DAP, all of the kernels had turned fully purple. The 25 DAP kernels were used for anthocyanin quantification at sweetcorn eating stage.

## 4. Discussion

### 4.1 Anthocyanin profiling of mature round starchy kernels

The current research addressed the anthocyanin profile of the novel purple-pericarp sweetcorn homozygous and heterozygous lines, and the hypothesis that the homozygous line (‘Tim1’) should produce a similar amount of anthocyanin present in its purple-pericarp ‘Costa Rica’ parent. It was observed that anthocyanin synthesis was affected by kernel maturity at harvest, with colour development increasing in conjunction with a progression of anthocyanin development across the kernel surface. Pigmentation was present in both the aleurone and pericarp tissues of the ‘Costa Rica’ parent and ‘Tim1’ developed line. Importantly, anthocyanin development occurred during the sweetcorn eating stage, and at 25 DAP close to the entire kernel surface was purple. The anthocyanin/ colour relationship was also affected by both total anthocyanin concentration and the ratio of cyanidin:pelargonidin.

From this current study, it was observed that the mature starchy ‘Costa Rica’ parental line had 97% purple pigmentation (cyanidin- and peonidin-based anthocyanin) and 3% red pigmentation (pelargonidin-based anthocyanin) pigments (Fig 2a). Cyanidin and peonidin together form a purple colour, with previous study reporting that peonidin is formed by methylation of cyanidin (Petrussa et al., 2013). By contrast, the purple shrunken kernels would have higher total anthocyanin than the purple round kernels, since there is no starch to dilute it.

Anthocyanin-biosynthesis dominant genes (at least one copy of each, e.g., *A1*) is required (Chaves-Silva et al., 2018) for pigmentation of anthocyanin in purple starchy maize. Moreover, study by Petroni et al., (2014) and Petroni and Tonelli, (2011) revealed that one transcription factor gene from Myb as well as bHLH transcription factor family is also required or the biosynthesis of anthocyanin. Earlier research by Procissi et al., (1997) demonstrated that purple pigment (light-independent) is formed in the maize pericarp in presence of *Pl1* (*purple plant1*), which is *a* Myb transcription factor allele. Recent studies by Chatham and Juvik, (2021; and Chatham et al., (2019) have shown that anthocyanin development in maize pericarp also requires bHLH transcription factor allele, *R1*.

The ‘Costa Rica’ purple maize parent produced the highest amount of anthocyanin, means that the *A1* (*Anthocyanin1*), *Pl1* (*Purple plant1*) and *R1* (*Coloured1*) genes are homozygous dominant in ‘Costa Rica’ parental line. Previous study by Chandler et al., (1989) indicated that in the pericarp of purple maize, anthocyanin biosynthesis is induced by the presence of anthocyanin biosynthesis dominant structural and transcription factor regulatory genes.

Aleurone pigmented kernels produced seven-fold less anthocyanin than the ‘Costa Rica’ genotype (Fig 2a). Previous research by Paulsmeyer et al., (2017) found that pericarp-coloured kernels had eight-times more anthocyanins in comparison to the aleurone-coloured kernels. This is why pericarp-pigmented maize is more economically valuable as a pigment source than aleurone-pigmented maize.

### 4.2 Anthocyanin profiling of mature shrunken non-starchy kernels

The F3 purple shrunken genotype, ‘Tim2’, had five-fold higher anthocyanin than the ‘Costa Rica’ parent (Fig 2b). The reason for this is largely that the ‘Costa Rica’ parent had round purple kernels with a starchy endosperm that diluted the amount of anthocyanin when compared to the ‘Tim2’ purple shrunken genotype, which has no starch in the endosperm, and, as a result, is shrunken with pericarp making up more of the mature kernel tissue. This finding agrees with an early research by Hong et al., (2020).

As expected with a non-pigmented sweetcorn, there was no anthocyanin present in the ‘Tims-white’ shrunken sweetcorn parent, which indicates that anthocyanin pigmentation structural gene was recessive (*a1*) in this parent. Earlier study by Chhabra et al., (2019) has also confirmed that a non-functional *a1* allele is responsible for the lack of anthocyanin in *shrunken2* sweetcorn.

### 4.3 Anthocyanin derivatives of sweetcorn eating stage kernels

No significant (P<0.005) difference was found between ‘Tim1’ and ‘Costa Rica’ for total anthocyanin concentration at 25 DAP (Fig 3c), however percentage of purple (cyanidin with peonidin) and red (pelargonidin) pigments between them were significantly different (Fig 3c). In ‘Tim1’, more pelargonidin is found compared to the ‘Costa Rica’ parent.

The *A1*, *Pl1* and *R1* genes are homozygous dominant in the developed F6 line ‘Tim1’ as its ‘Costa Rica’ parent. Therefore, it is likely that a change in the *Pr1* (*purple/red aleurone1*) gene is responsible for more pelargonidin in ‘Tim1’ than ‘Costa Rica’, which indicates that a heterozygous (*Pr1pr1*) or homozygous recessive (*pr1pr1*) allele is present in ‘Tim1’ and is the cause of increased proportion of pelargonidin compared to ‘Costa Rica’ parent.

Earlier research by Sharma et al., (2011) showed that for dominant *Pr1* alleles (*Pr1Pr1*), purple colour is produced due to cyanidin-based anthocyanin accumulation; whereas recessive *pr1* alleles (*pr1pr1*) is responsible for red colour due to the accumulation of pelargonidin-based anthocyanin; with the heterozygous form of *Pr1* alleles (*Pr1pr1*) yielding an intermediate reddish-purple colour. As the kernels of the F6 line, ‘Tim1’ were reddish-purple, this may indicate that the *Pr1* gene may be heterozygous in this line.

### 4.4 Colour measurement

Colour was measured objectively (McGuire, 1992) to observe the relationship between total anthocyanin concentration with either hue or chroma values (Fig 4). The starchy round kernels of ‘Costa Rica’ showed significantly (P<0.005) lower H* value (13.9) (Fig 4c), corresponding to a darker colour. Previous study by Hernandez et al., (2004) found H* value ranged from 9.5 to 34.2 in different purple Peruvian maize lines.

Within the developed F3 lines, ‘Tim2’ had a significantly lower (P<0.005) H* value than other shrunken kernels and total highest anthocyanin in comparison to them (Fig 4c). This means that the lines with lower hue angle had higher total anthocyanin concentrations, as observed in round ‘Costa Rica’ above, as well as shrunken F3 line, ‘Tim2’. No significant (P<0.005) difference was observed regarding H* value between the ‘Tim1’ and ‘Tim4’ lines (Fig 4c), which was also supported by the similar anthocyanin profiles in each of these (Fig 2b).

### 4.5 Anatomy of kernel anthocyanin development

Anthocyanin development was only apparent in the pericarp, aleurone, as well as vitreous endosperm, which would have been derived from the aleurone layer. The presence of anthocyanin in the vitreous endosperm was also reported earlier by Cortés et al., (2006). However, there was no anthocyanin developed in the embryo or starchy endosperm (Fig 5a). It indicated that the genes responsible for anthocyanin development were responsible for anthocyanin development only in the pericarp and/or aleurone layers, and not in the embryo or starchy endosperm.

The anatomical study also revealed that due to the presence of starch in the endosperm, the round purple kernels produced less anthocyanin than the shrunken purple kernels (Fig 2a). In addition, significantly less anthocyanin was produced in ‘Tims-aleurone’ (27.65 mg/100g FW) kernels than in the ‘Costa Rica’ (194.47 mg/100g FW) purple-pericarp kernels. The reason behind this is that the maternal pericarp tissue consists of several outer layers, by contrast, non-maternal aleurone remains below the pericarp (Fig 5a) and consists of single layer of cell (Becraft and Yi, 2011). Earlier research by Luna-Vital et al., (2017) has also found significantly higher amount of anthocyanin in pericarp-pigmented corn than aleurone-pigmented corn.

### 4.6 Physiology of kernel anthocyanin development

During the physiological development of the kernel, it was observed that with increasing kernel maturity, the intensity of pericarp colour also increased (Fig 5b). Previous research by Hong et al., (2020) demonstrated that anthocyanin spreads across the pericarp with increasing maturity. Therefore, it can be assumed that more anthocyanin is produced with increasing DAP. This finding is in agreement with a previous study of purple sweetcorn based on the *brittle1* mutation (Hong et al., 2020) and purple maize (Li et al., 2019), both of which demonstrated that anthocyanin development increased with kernel maturity.

Anthocyanin has different health benefits including the prevention of hypertension as well as different types of cancers. Total anthocyanin found in red currants, red plums and coloured strawberries were 12.8 mg/100g FW, 30.1 mg/100g FW and 60 mg/100g FW, respectively (Aaby et al., 2012, Proteggente et al., 2002, Wu et al., 2006). At eating stage (25 DAP) of current study, the developed F6 line ‘Tim1’ produced 253.03 mg/100g FW total anthocyanin, which is significantly much higher compared to the anthocyanin found in the above-mentioned pigmented crops.

## 5. Conclusion

Novel purple-pericarp super-sweetcorn lines were assessed for their anthocyanin concentration both at sweetcorn-eating and mature stages. The ‘Tim1’ line developed a similar total anthocyanin concentration to the ‘Costa Rica’ parent when assessed at sweetcorn-eating stage, while at full maturity, it actually surpassed the purple maize parent, but this was mainly due to the presence of non-pigmented starch diluting the total anthocyanin of the latter. On the other hand, there is a lack of anthocyanin in standard white/yellow sweetcorn. Therefore, the high anthocyanin concentration found in the developed purple sweetcorn ‘Tim’ lines of current study could potentially be used for preventing non-contaminated diseases.

